# CBIcall: a configuration-driven framework for variant calling in large sequencing cohorts

**DOI:** 10.64898/2026.03.23.713646

**Authors:** Manuel Rueda, Dietmar Fernandez-Orth, Ivo G. Gut

## Abstract

**Motivation:** Variant calling for next-generation sequencing (NGS) data relies on a diverse ecosystem of tools and workflows. Large-scale collaborative studies increasingly adopt federated analysis, where each institution processes sensitive data locally using standardized pipelines. Deploying identical pipelines across multiple centers remains challenging because heterogeneous software environments and computing policies can cause workflow divergence and inconsistent results.

**Results:** We developed CBIcall, a workflow-agnostic, configuration-driven framework that runs standardized variant-calling pipelines from raw FASTQ files to analysis-ready VCFs using a single YAML file. An execution driver validates user parameters, enforces compatibility across pipelines, analysis modes, workflow backends, genome builds, and tool versions, and records structured provenance for each run, ensuring consistent and reproducible pipeline execution across computing environments. CBIcall dispatches validated workflows through Bash or Snakemake backends and provides production-ready pipelines for germline WES, WGS (single-sample or cohort joint genotyping following GATK Best Practices), and mitochondrial DNA analysis. We validated CBIcall on public datasets and deployed it in the EU HEREDITARY project, processing 1,111 samples with both WES and mtDNA pipelines on an institutional HPC system, demonstrating its suitability for reproducible cohort-scale genomic analyses.

**Availability and implementation:** CBIcall is open source (GPLv3) and distributed with ready-to-run pipelines; full dependency and installation documentation is available at https://github.com/CNAG-Biomedical-Informatics/cbicall.

## 1 Introduction

Next-generation sequencing (NGS), introduced in 2005, has transformed genomic research by making DNA and RNA profiling scalable and cost-efficient (Metzker 2010, Goodwin, McPherson, and McCombie 2016). As sequencing technologies have matured, the software ecosystem has evolved from simple command-line utilities to complex workflow engines and multi-stage analysis (Köster and Rahmann 2012, Amstutz *et al*. 2016, Di Tommaso *et al*. 2017, Galaxy Community 2024). This diversity provides analytical flexibility, but it also creates practical challenges for collaborative studies that must analyze large cohorts using stable, consistent methods across sites.

Many European research projects now use federated analysis models, where each institution processes genomic data locally to comply with legal and ethical requirements (Rehm *et al*. 2021). A key challenge is executing identical, validated pipelines across heterogeneous HPC environments that differ in software stacks, scheduling policies, and file system conventions. Although many publicly available workflows are analytically robust, they are frequently distributed as workflow templates without a standardized validation layer that enforces configuration correctness, compatible tool versions, and consistent runtime environments. As a result, deployments across different institutions often require site-specific wrappers and manual adjustments, which may introduce workflow divergence and compromise reproducibility (Leipzig 2017, Grüning *et al*. 2018, Ahmed *et al*. 2021).

To address these challenges, we developed CBIcall within the EU Horizon Europe HEREDITARY project (https://hereditary-project.eu). The framework was developed in the context of large collaborative sequencing projects at the Centro Nacional de Análisis Genómico (CNAG), where robust and reproducible execution of standardized pipelines across heterogeneous computing environments is required. Rather than introducing a new workflow engine, CBIcall provides a configuration-driven validation and execution layer above existing workflow backends such as the Linux Bash shell (Ramey and Fox 2025) and Snakemake (Köster and Rahmann 2012). It enables validated nuclear (McKenna *et al*. 2010) and mitochondrial variant-calling pipelines (Calabrese *et al*. 2014) to run “out of the box” from a single YAML file while enforcing compatibility rules and recording structured provenance for each run.

## 2 Methods

### 2.1 Overall design and system requirements

CBIcall is a configuration-driven framework designed to ensure consistent and reproducible variant-calling workflows across institutional computing environments, including high-performance computing (HPC) clusters commonly used in large-scale sequencing centers. The system provides ready-to-use pipelines for nuclear and mitochondrial DNA variant calling, processing paired-end FASTQ files into analysis-ready VCF files.

CBIcall adopts a two-layer configuration model:

1. User parameters YAML: specifies the analysis intent, including input samples, pipeline and mode selection, genome build, and tool parameters (example provided in Sup. Text S1).
2. Workflow registry YAML: maps pipeline definitions and workflow backends to executable workflow scripts (Bash or Snakemake), optionally referencing shared workflow components (example provided in Sup. Text S2).

Separating these layers minimizes configuration drift and prevents workflow divergence across different runs and installations.

### 2.2 Execution control layer

The CBIcall driver (implemented in Python 3) serves as the framework’s central execution controller (Figure 1). For each run, the driver loads the user’s YAML parameters and validates all input parameters against controlled vocabularies. It enforces compatibility rules across analysis modes, genome builds, workflow backends, and tool versions (including specific version constraints for GATK (McKenna *et al*. 2010)). Workflow definitions are validated using JSON Schema Draft 2020-12. The system then creates a deterministic project directory structure for each execution and launches the workflow using the appropriate backend.

**Figure 1.**
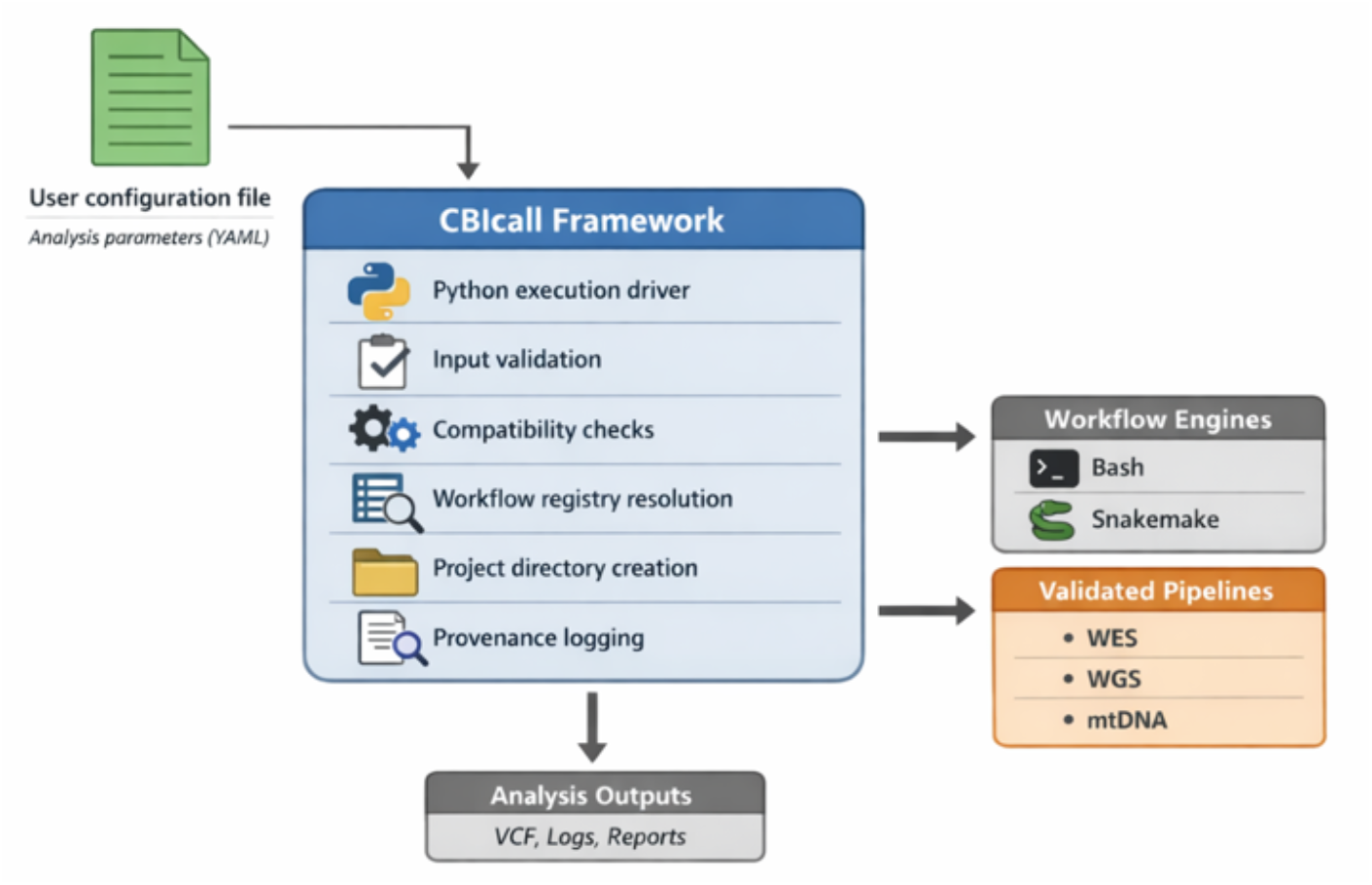
Overview of the CBIcall execution model. Users define analysis parameters in a YAML configuration file. The CBIcall execution driver validates inputs, enforces compatibility rules, resolves the requested pipeline via a workflow registry, and launches the workflow using Bash or Snakemake to generate variant-calling outputs.

During execution, the system records structured metadata—including software versions, analysis parameters, runtime context, and task-level execution provenance—in a JSON file (default: ‘log.json’; example provided in Sup. Text S3). This supports both auditing and reproducible reanalysis.

### 2.3 Workflow backends and validated pipelines

CBIcall currently supports two workflow backends: Bash and Snakemake. The Bash backend includes ready-to-use pipelines for whole-exome sequencing (WES), whole-genome sequencing (WGS), and mitochondrial DNA (mtDNA) variant calling. These support both single-sample and cohort-level processing, using GATK 3.5 (McKenna *et al*. 2010) for legacy datasets following previously described workflows (Rueda *et al*. 2017) and GATK 4.6 for current analyses (Poplin *et al*. 2017). Nuclear germline pipelines follow GATK Best Practices (DePristo *et al*. 2011, Van der Auwera *et al*. 2013) and include read alignment with BWA-MEM (Li 2013), coordinate sorting, duplicate marking, base quality score recalibration, persample gVCF generation using *HaplotypeCaller*, cohort-level genotyping with *GenotypeGVCFs*, variant quality score recalibration, and standard post-processing filters (see online documentation).

The Snakemake backend is provided as a validated reference implementation and supports WES and WGS pipelines using GATK 4.6. In principle, users can integrate external Snakemake workflows—including those from community repositories such as WorkflowHub (Gustafsson *et al*. 2025) or Dockstore (Yuen *et al*. 2021)—by registering them in the YAML workflow registry. These workflows execute within CBIcall’s runtime directory, which enforces a consistent project structure and logging. Validation and execution logic remain implemented in the core framework. The architecture is extensible and would allow additional workflow engines (e.g., Nextflow) to be integrated using the same execution model.

For mitochondrial DNA, variant calling uses MToolBox (which requires GATK 3.5) (Calabrese *et al*. 2014). Mitochondrial DNA analyses produce a VCF file, a prioritized annotation file, and an interactive HTML report (see Sup. Figure S1) adapted from a previously described framework (Rueda and Torkamani 2017). For WES and WGS analyses, we provide annotation, visualization, and Beacon v2 API conversion through the *beacon2-cbi-tools* software (formerly *beacon2-ri-tools*), described elsewhere (Rueda *et al*. 2022).

### 2.4 Containerization and portability

CBIcall is distributed as a container image that includes the execution framework and its Python dependencies, and it can alternatively be installed from source. External resources—including reference genomes, annotation databases, and third-party tools such as GATK, BWA-MEM, and Samtools (see list at Sup. Text S4)—are distributed separately and must be downloaded and placed in a designated resource directory as described in the online documentation.

In containerized environments, users mount this external resource directory into the container and register it through a configuration file. On institutional HPC systems that do not support Docker, the same container image can be executed using Apptainer (formerly Singularity), enabling consistent deployment across heterogeneous computing environments.

## 3 Results: Use cases and validation

### 3.1 Study cohorts and sequencing data

Because CBIcall operates as a validation and execution layer above existing workflow backends, its performance is best evaluated through realistic end-to-end analyses rather than isolated benchmarking.

To illustrate the robustness and flexibility of the framework, we applied CBIcall to two representative use cases using real sequencing datasets processed within an institutional (CNAG) HPC environment. To evaluate CBIcall in a realistic collaborative cohort setting, we analyzed an initial dataset of 618 WES samples from the National Institute of Neurological Disorders and Stroke (NINDS) Parkinson’s Disease study (dbGaP accession phs001172.v1.p2). After quality control and population structure assessment, 10 samples were excluded due to ethnic stratification (see Sup. Figure S2), resulting in a final dataset of 608 samples. These were analyzed as part of the HEREDITARY-CNAG Parkinson’s cohort, together with 503 control samples from the 1000 Genomes (1000G) Project, a predominantly WGS resource that also includes WES data (Genomes Project *et al*. 2015), forming a study cohort of 1,111 samples. Both nuclear and mitochondrial analyses described below were performed on this same WES-derived cohort.

### 3.2 Use case 1: Nuclear variant calling in a large WES cohort

To demonstrate large-scale nuclear variant calling in this cohort, starting from FASTQ files, we processed all samples using the CBIcall WES pipeline with the b37 (hg19) reference genome (selected for compatibility with validated GATK resource bundles), followed by standard *bcftools* (Danecek *et al*. 2021) post-filtering. This produced two QC-matched callsets that differed primarily in their genotyping strategy while sharing identical alignment and preprocessing steps, enabling a direct comparison of single-sample and cohort-level calling approaches:

i. Single-sample mode: We called each sample individually using GATK *HaplotypeCaller*, producing per-sample gVCF and filtered VCF files. We then merged the VCFs (*bcftools merge*) without cohort-level re-genotyping.
ii. Cohort-joint genotyping mode: The 1,111 gVCF files generated in single-sample mode were combined and jointly genotyped using GATK *GenotypeGVCFs*. The resulting VCF was then subjected to the same filtering procedure applied in the single-sample analysis.

As can be seen in Figure 2A, there was a strong intersection of variants in the VCF coming from single-sample mode and the cohort-genotyping mode. After applying a PASS filter (Figure 2B), the joint-genotyped callset retained more variants that were filtered out in the merged single-sample callset, consistent with the expected advantages of cohort-level joint genotyping (see also Sup. Table S1).

**Figure 2.**
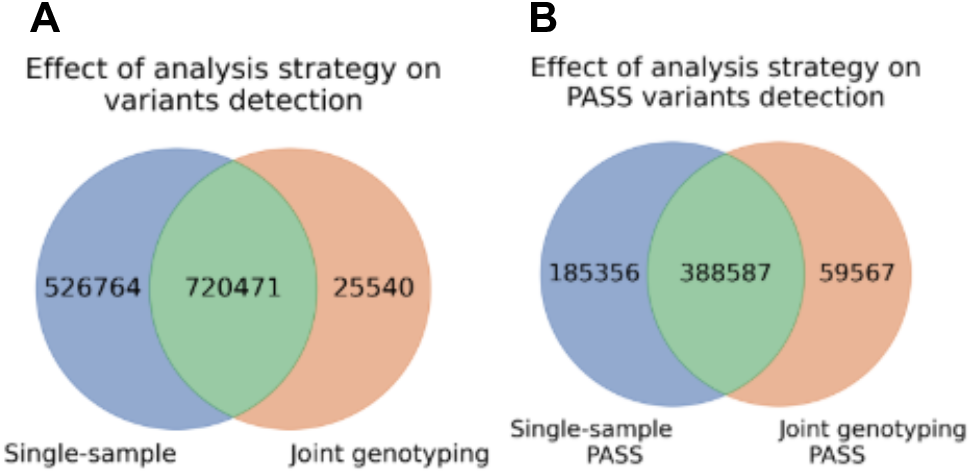
Configuration-consistent comparison of genotyping strategies using CBIcall. Venn diagram showing the overlap between variant sets obtained from single-sample variant calling (followed by VCF merging), and from cohort-level joint genotyping of the same 1,111 samples. A) Variants obtained directly from calling. B) Variants remaining after PASS filter.

The comparability of the phs001172 and 1000G cohorts was evaluated using per-sample variant counts (see Sup. Table S2) and sequencing depth (DP; see Sup. Figure S3) as quality control metrics to identify potential technical biases prior to integration. Distributions of WES-derived SNVs and INDELs were highly similar across cohorts, supporting their suitability for comparative downstream analyses. Variant counts obtained under the different genotyping strategies were consistent, indicating no systematic shifts in variant detection. Likewise, DP distributions across all samples, including cases and controls processed under both analytical modes and mitochondrial analysis, were broadly comparable, further confirming the absence of method-specific bias.

Finally, we performed a Principal Component Analysis (PCA) based on sample genotypes to assess potential population structure or batch effects. Controls and cases displayed a largely overlapping distribution with no clear group separation, indicating that the integrated dataset does not exhibit major systematic differences and is suitable for downstream analyses (see Sup. Figure S4).

### 3.3 Use case 2: Mitochondrial variant calling from WES data

In this second use case, mitochondrial genome analysis was performed on the combined study cohort (1,111 WES samples) using MToolBox as part of the CBIcall analysis workflow. Variant calling was successful in approximately 95% of samples. Overall, WES provided sufficient mtDNA coverage for the vast majority of samples (see Sup. Figure S3). In a limited number of cases (10/503 for 1000G, 1.98%, and 55/608 for dbGaP, 9.05%), coverage was below MToolbox’s analytical threshold required for reliable variant calling. This reflects the expected off-target nature of mitochondrial read capture in WES (Calabrese *et al*. 2014, Griffin *et al*. 2014, Poole *et al*. 2021).

We performed two analyses on the annotated variants. First, heteroplasmic variants (defined as heteroplasmic fraction > 0.3) were grouped by genomic locus, and their cohort-specific and relative frequencies (in percentage) were assessed for cases and controls (Figure 3). The results show that despite the different sequencing strategies on both source cohorts, only minor differences were observed in the distribution of heteroplasmic variants across loci (Diroma *et al*. 2014, Rueda and Torkamani 2017, Laricchia *et al*. 2022).

**Figure 3.**
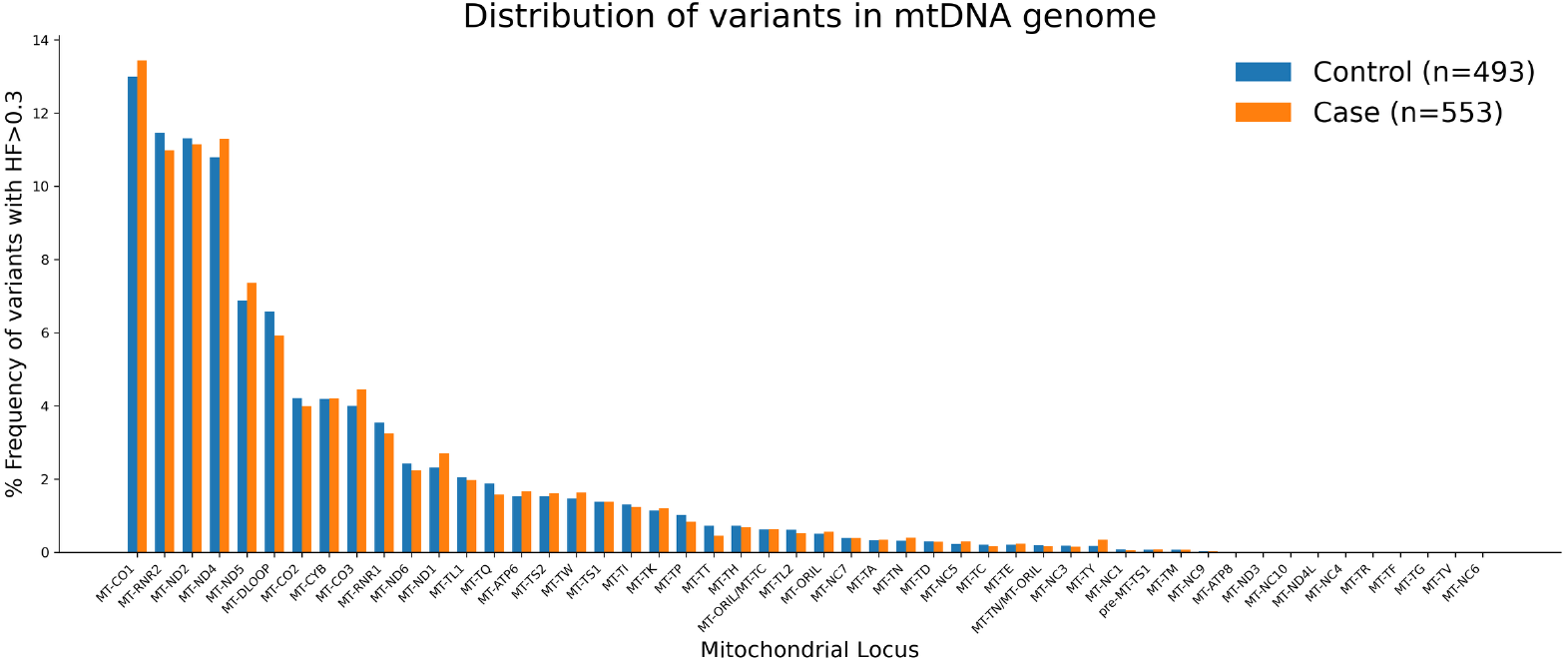
Bar plot showing the distribution of heteroplasmic variants across the 1,111 control and case samples. A heteroplasmy threshold of 0.3 was applied, and mitochondrial loci were sorted in descending order according to the number of variants.

Finally, to assess whether heteroplasmic variants were uniformly distributed across the mitochondrial genome, we generated a scatterplot in which loci were ordered according to their length (bp). As expected, the number of heteroplasmic variants per locus showed a consistent increasing trend with locus size in both case and control cohorts (see also Sup. Figure S5) (Diroma *et al*. 2014, Rueda and Torkamani 2017, Laricchia *et al*. 2022).

## Acknowledgements

Not available.

## Supplementary data

Supplementary data are available at *Bioinformatics Advances* online.

## Conflict of interest

Declarations. Ethics approval and consent to participate: A data access request was submitted to the NCBI Database of Genotypes and Phenotypes (dbGaP) to obtain the FASTQ files for National Institute of Neurological Disorders and Stroke (NINDS) Exome Sequencing in Parkinson (dbGaP; accession no. phs001172.v1.p2; approval 142786-2).

Consent for publication: Not applicable.

Competing interests: The authors declare no competing interests.

## Data availability

The data underlying this article are available in 1000 Genomes project at https://www.internationalgenome.org/data-portal/data-collection/phase-3 and dbGaP at https://dbgap.ncbi.nlm.nih.gov/beta/study/phs001172.v1.p2/.

