## Supplementary Data for "CBIcall: a configuration-driven framework for variant calling in large sequencing cohorts"

#### Supplementary Data Contents:

##### Supplementary Methods

**Text S1: Example CBIcall user configuration**

This example YAML file illustrates a typical user configuration used to run a whole-exome sequencing (WES) analysis with CBIcall. The configuration specifies the mode ("single" or "cohort"), selected analysis pipeline, workflow backend, input sample, genome build, and other analysis parameters.

Example configuration excerpt:

```
---
mode:            single
pipeline:        wes
workflow_engine: bash
gatk_version:    gatk-4.6
input_dir:       CNAG999_exome/CNAG99901P_ex
genome:          b37
cleanup_bam:     false
```

**Text S2: Workflow registry excerpt**

This excerpt from the CBIcall workflow registry illustrates how pipeline definitions are resolved into backend-specific executable workflows. The registry maps analytical pipelines defined in the user configuration to implementation-specific scripts (e.g., Bash workflows), while preserving a stable user-facing configuration interface.

Example registry structure:

```
---
workflows:
  bash:
    base_dir: "workflows/bash"
    versions:
      gatk-4.6:
        common:
          env: "env.sh"
          coverage: "coverage.sh"
          jaccard: "jaccard.sh"
          vcf2sex: "vcf2sex.sh"
        pipelines:
          wes:
            single: "wes_single.sh"
            cohort: "wes_cohort.sh"
          wgs:
            single: "wgs_single.sh"
            cohort: "wgs_cohort.sh"
```

**Text S3: Execution provenance record**

This excerpt from a CBIcall execution provenance file (log.json) illustrates the structured metadata recorded for a representative analysis run. The record captures software versions, pipeline and workflow identifiers, and runtime parameters automatically generated during execution. These provenance records support auditing and reproducible reanalysis across computing environments.

Example provenance record excerpt:

```
{
  "arg": {
    "debug": null,
    "help": false,
    "man": false,
    "nocolor": false,
    "paramfile": "param.yaml",
    "show_version": false,
    "threads": 6,
    "verbose": false
  },
  "config": {
    "allow_partial_run": false,
    "arch": "arm64",
    "capture_label": "GATK_bundle_b37",
    "compression_cmd": "/usr/bin/pigz -p 5",
    "date": "Mon Mar 23 11:52:42 2026",
    "gatk_version": "gatk-4.6",
    "genome": "b37",
    "host_threads": 6,
    "host_threads_minus_one": 5,
    "hostname": "mrueda-ws1",
    "inputs": {
      "input_dir":
"/media/mrueda/2TBS/CNAG/Project_CBI_Call/cbicall/examples/input/CNAG999_exome/CNAG99901P_ex",
      "sample_map": null
    },
    "mode": "single",
    "output_basename": "CNAG99901P_ex",
    "pipeline": "wes",
    "project_dir":
"/media/mrueda/2TBS/CNAG/Project_CBI_Call/cbicall/examples/input/CNAG999_exome/CNAG99901P_ex/cbica
ll_bash_wes_single_b37_gatk-4.6_177426316206646",
    "run_id": "177426316206646",
    "run_mode": "full",
    "user": "mrueda",
    "version": "1.0.0",
    "workflow": {
      "config_file": null,
      "engine": "bash",
      "entrypoint":         "/media/mrueda/2TBS/CNAG/Project_CBI_Call/cbicall/workflows/bash/gatk-
4.6/wes_single.sh",
      "gatk_version": "gatk-4.6",
      "helpers": {
        "coverage":         "/media/mrueda/2TBS/CNAG/Project_CBI_Call/cbicall/workflows/bash/gatk-
4.6/coverage.sh",
        "env": "/media/mrueda/2TBS/CNAG/Project_CBI_Call/cbicall/workflows/bash/gatk-4.6/env.sh",
        "jaccard":          "/media/mrueda/2TBS/CNAG/Project_CBI_Call/cbicall/workflows/bash/gatk-
4.6/jaccard.sh",
        "vcf2sex":          "/media/mrueda/2TBS/CNAG/Project_CBI_Call/cbicall/workflows/bash/gatk-
4.6/vcf2sex.sh"
      },
      "mode": "single",
```

```
      "pipeline": "wes"
    },
    "workflow_engine": "bash",
    "workflow_rule": null
  },
  "param": {
    "allow_partial_run": false,
    "cleanup_bam": false,
    "gatk_version": "gatk-4.6",
    "genome": "b37",
    "input_dir":
"/media/mrueda/2TBS/CNAG/Project_CBI_Call/cbicall/examples/input/CNAG999_exome/CNAG99901P_ex",
    "mode": "single",
    "organism": "Homo sapiens",
    "output_basename": null,
    "pipeline": "wes",
    "project_dir": "cbicall",
    "sample_map": null,
    "technology": "Illumina HiSeq",
    "workflow_engine": "bash",
    "workflow_rule": null
  }
}
```

#### Text S4: External databases and third-party utilities distributed with the CBIcall resource package

| Category | Resource | Version / Build | Archive path | Purpose | Reference |
|---|---|---|---|---|---|
| Reference genome | Human genome | hg38 | Databases/genomes/hg38 | Reference genome used for alignment and variant calling | Genome Reference Consortium |
| Annotation database | dbSNP | b144 (GRCh37) | Databases/dbSNP/human_9606_b144_GRCh37p13 | Known variants used for annotation and recalibration | (Sherry *et al.* 2001) |
| Annotation database | dbSNP | b146 (GRCh38) | Databases/dbSNP/human_9606_b146_GRCh38p2 | Known variants annotation and recalibration | (Sherry *et al.* 2001) |
| Workflow resources | GATK resource bundle | — | Databases/GATK_bundle | Known-sites resources required by GATK workflows | Broad Institute |
| Capture resources | Agilent SureSelect | — | Databases/Agilent_SureSelect | Target interval definitions used for exome sequencing analyses | Agilent Technologies |
| Mitochondrial resources | mtDNA references | — | Databases/mtDNA | Mitochondrial reference sequences (rCRS and RSRS) and derived hg19-based FASTA files (chrM, hg19RCRS, hg19RSRS), compressed FASTA resources, and GMAP alignment databases used by mitochondrial workflows | (Calabrese *et al.* 2014) |
| Alignment software | BWA | 0.7.18 | NGSutils/bwa-0.7.18* | Read alignment to the reference genome | (Li Heng and Durbin 2009, Li Heng 2013) |
| Variant analysis framework | GATK | 3.5; 4.6 | NGSutils/gatk/ | Germline variant calling and preprocessing workflows | (McKenna *et al.* 2010) |
| BAM processing utilities | SAMtools | 0.1.19; 1.3 | NGSutils/samtools-* | Manipulation of BAM/SAM alignment files | (Li H. *et al.* 2009) |
| BAM processing utilities | Picard | 2.25 | NGSutils/picard-2.25 | Duplicate marking and BAM processing | Broad Institute |
| Genomic interval utilities | BEDTools | 2.x | NGSutils/bedtools2* | Genomic interval and feature manipulation | (Quinlan and Hall 2010) |
| Runtime dependency | Java | 8 | NGSutils/java8 | Java runtime required by GATK and Picard | Oracle |
| Runtime dependency | Python | 2.7 (portable) | NGSutils/python_2.7/linux-x86_64/python27_portable | Legacy runtime required by specific pipeline components | Python Software Foundation |
| mtDNA analysis pipeline | MToolBox | 1 | NGSutils/MToolBox-master | Pipeline for mitochondrial genome assembly and heteroplasmy analysis | (Calabrese *et al.* 2014) |

* Indicates directories containing architecture-specific builds (e.g., x86_64 and ARM64).

CBIcall v1.0.0 includes a curated resource package containing reference datasets and third-party bioinformatics utilities required by the validated workflows. The main components of this package are summarized in the Table above.

### Figure S1: Example mtDNA HTML report

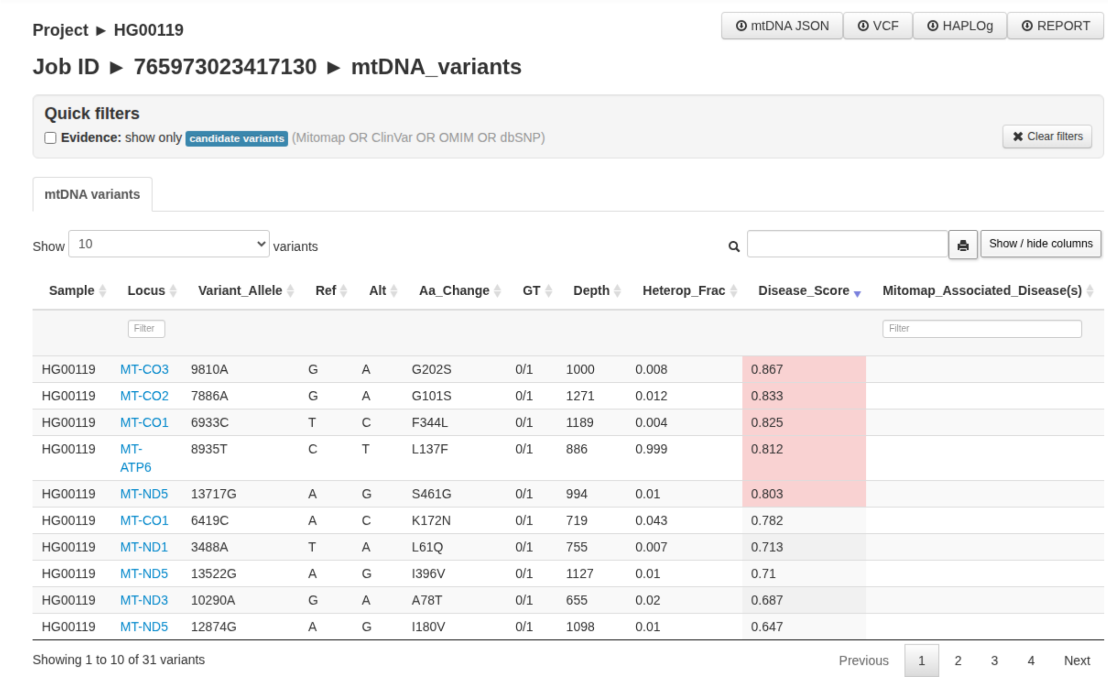

Project ► HG00119

mtDNA JSON | VCF | HAPLOg | REPORT

Job ID ► 765973023417130 ► mtDNA_variants

Quick filters

Evidence: show only candidate variants (Mitomap OR ClinVar OR OMIM OR dbSNP)

Clear filters

mtDNA variants

Show 10 variants

Show / hide columns

| Sample | Locus | Variant_Allele | Ref | Alt | Aa_Change | GT | Depth | Heterop_Frac | Disease_Score | Mitomap_Associated_Disease(s) |
|---|---|---|---|---|---|---|---|---|---|---|
| HG00119 | MT-CO3 | 9810A | G | A | G202S | 0/1 | 1000 | 0.008 | 0.867 | |
| HG00119 | MT-CO2 | 7886A | G | A | G101S | 0/1 | 1271 | 0.012 | 0.833 | |
| HG00119 | MT-CO1 | 6933C | T | C | F344L | 0/1 | 1189 | 0.004 | 0.825 | |
| HG00119 | MT-ATP6 | 8935T | C | T | L137F | 0/1 | 886 | 0.999 | 0.812 | |
| HG00119 | MT-ND5 | 13717G | A | G | S461G | 0/1 | 994 | 0.01 | 0.803 | |
| HG00119 | MT-CO1 | 6419C | A | C | K172N | 0/1 | 719 | 0.043 | 0.782 | |
| HG00119 | MT-ND1 | 3488A | T | A | L61Q | 0/1 | 755 | 0.007 | 0.713 | |
| HG00119 | MT-ND5 | 13522G | A | G | I396V | 0/1 | 1127 | 0.01 | 0.71 | |
| HG00119 | MT-ND3 | 10290A | G | A | A78T | 0/1 | 655 | 0.02 | 0.687 | |
| HG00119 | MT-ND5 | 12874G | A | G | I180V | 0/1 | 1098 | 0.01 | 0.647 | |

Showing 1 to 10 of 31 variants

Previous 1 2 3 4 Next

Downloadable files:

- **mtDNA JSON** A JSON file with the results from `mit_prioritized_variants.txt`.
- **Report:** A tsv file including all the annotations for each variant. Name of the file `mit_prioritized_variants.txt`.
- **Haplog:** A tsv file including the predicted haplogroup for each sample. Name of the file `mt_classification_best_results.csv`.
- **VCF:** A text file consisting of all the variants in the VCF format. Name of the file `VCF_file.vcf`.

HTML table:

In this tab SG-ADVISER mtDNA displays a browsable table consisting of the most relevant fields relative to the variant annotation:

- **Sample**: The full name of each sample.
- **Locus**: The location on the mitochondrial chromosome.
- **Variant_Allele**: The position in the mitochondrial chromosome + the alternative allele format.
- **Ref**: The reference allele (mitochondrial reference genome: RSRS).
- **Alt**: The alternative allele(s).
- **Aa_change**: The amino acid change if the variant falls in a coding region.
- **GT**: Genotype. 0:Ref, ≥1:Alt(s).
- **Depth**: The number of times this position is covered by reads.
- **Heterop_Frac**: The heteroplasmic fraction. Note that the confidence interval can be retrieved from the downloadable VCF file.
- **Other**: For other fields please consult MToolBox's manual.

- **Figure S2: dbGaP samples selection**

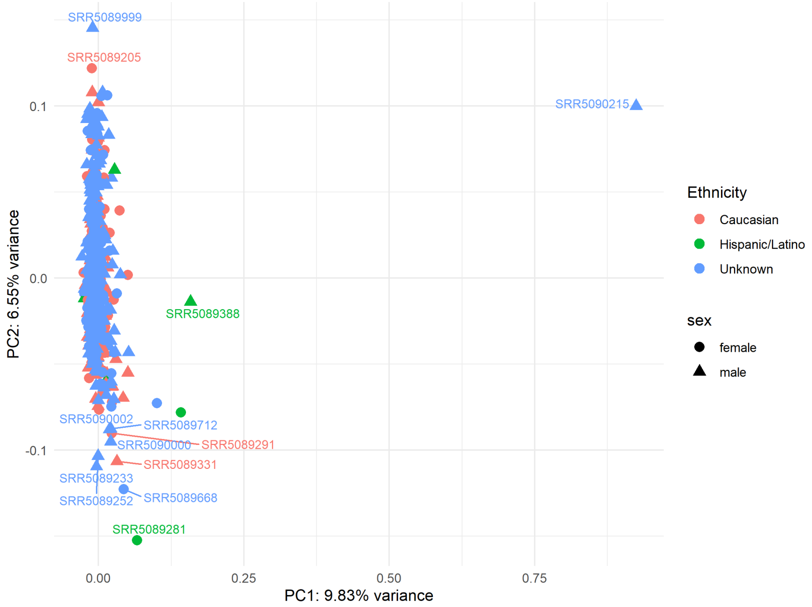

**Figure S2**. Principal component analysis (PCA) based on genotypes derived from CBIcall variant calls and converted to PLINK format for the 618 samples from the dbGaP dataset phs001172.v1.p2. Ten samples (Hispanic/Latino individuals and sample SRR5090215) clustered separately due to population stratification and were removed prior to downstream analyses.

For the CBIcall use cases, the selected dbGaP dataset (accession **phs001172.v1.p2**) comprised 618 samples. Population structure was assessed using principal component analysis (PCA), which identified 10 samples clustering separately due to ethnic stratification. These samples were excluded to avoid population structure bias in downstream analyses, resulting in a final dataset of 608 samples.

**Figure S3: Distribution of sequencing depth (DP) across analysis strategies.**

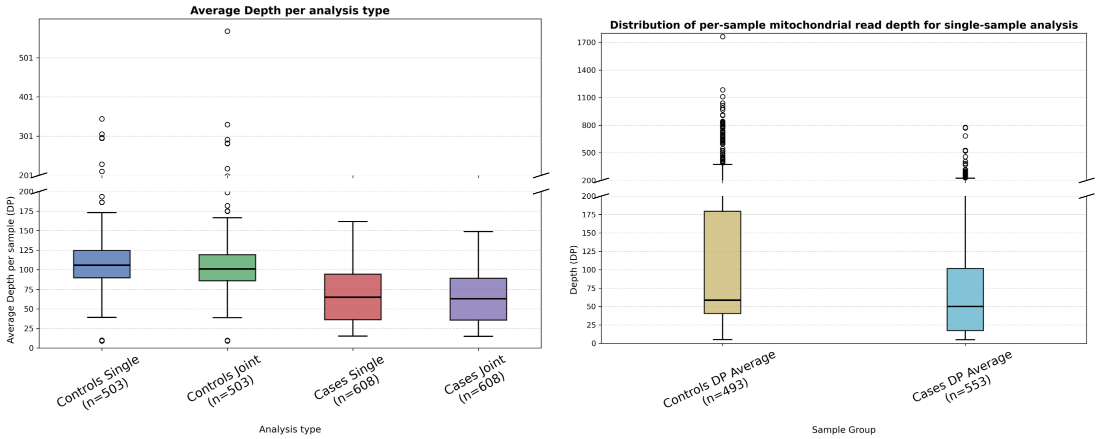

**Figure S3**. **Left**. Distribution of average sequencing depth (DP) per sample, grouped by analysis type for single vs joint analysis. **Right**. Distribution of average sequencing depth for mitochondrial single-sample analysis. A broken y-axis is applied to enhance visualization of inter-group variability (0–200, step 25; 200–600, step 100, left; 200–1700, step 300, right). Boxplots show that DP values are comparable across analysis strategies, indicating consistent per-sample coverage and that DP is greater in Control samples.

Boxplots of average sequencing depth (DP) per sample across analysis strategies are shown in Figure S3. For WES analyses, depth distributions were highly comparable between single-sample and joint genotyping modes. The 1000 Genomes cohort showed a median coverage of approximately 100× (106× for single-sample analysis and 101× for joint calling), whereas the phs001172 cohort showed a median coverage of approximately 64× (65× for single-sample analysis and 63× for joint calling).

For mitochondrial DNA analysis, median coverage was 59× for controls and 50× for cases. Thus, sequencing depth fell within the range generally considered sufficient for reliable variant detection.

**Figure S4: Principal component analysis (PCA) of cases and controls.**

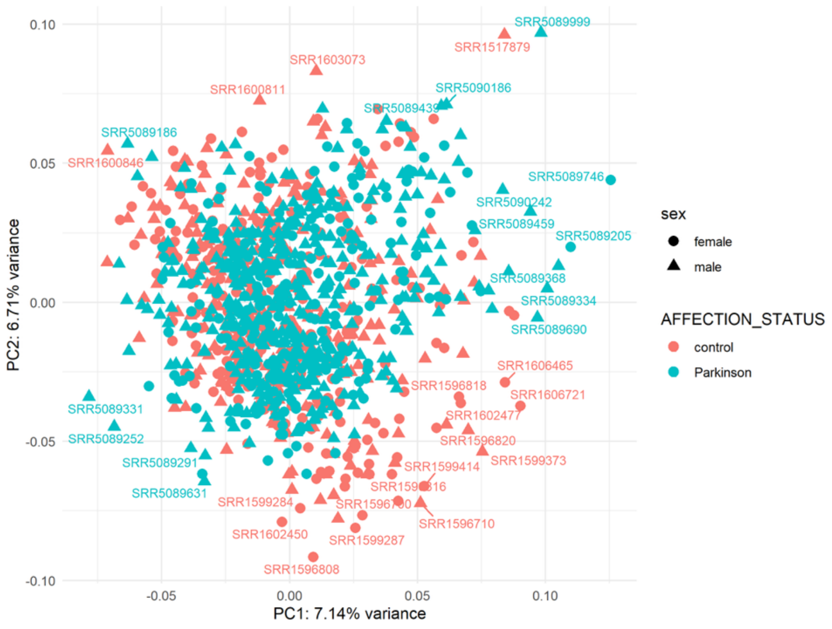

**Figure S4**. Principal component analysis (PCA) of genotypes derived from CBIcall variant calls and converted to PLINK format, using the intersection of variants between cases (dbGaP phs001172.v1.p2, n=608) and 1000 Genomes controls (n=503).

Principal component analysis (PCA) based on genotypes obtained from the single-sample calling analysis showed no clear separation between case and control samples. This result is consistent with expectations, as both cohorts were processed using comparable sequencing and variant-calling pipelines and therefore do not exhibit detectable technical batch effects.

**Figure S5: Relationship between mitochondrial locus length and heteroplasmic variant frequency.**

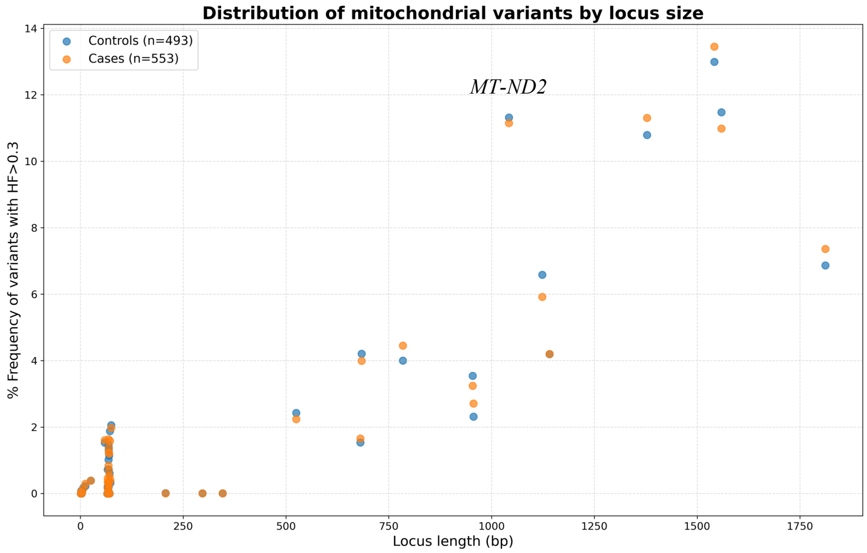

**Figure S5.** Scatter plot showing the relationship between mitochondrial locus length and the % frequency of heteroplasmic variants (HF > 0.3) across the case and control cohorts.

A detailed analysis of heteroplasmic variant distribution across mitochondrial loci revealed consistent increasing trend with locus size, consistent with previous observations.

Despite differences in sequencing strategies between cohorts, both case and control datasets exhibited similar patterns of heteroplasmic variation. A small number of short mitochondrial loci showed relatively higher variability in heteroplasmic variant counts. *MT-ND2* also displayed an increased number of variants. An explanation for these deviations could be influenced by technical factors such as variable mitochondrial coverage in whole-exome sequencing data and increased stochastic effects in short genomic intervals.

#### Table S1. Variant filtering statistics for single-sample and joint genotyping analyses

**Table S1.** Summary of variant counts before and after quality filtering (FILTER=PASS) in single-sample and joint genotyping analyses. The table reports the number of variants before filtering, the number retained after filtering, and the absolute and relative number of variants removed. Percentages indicate the proportion of variants discarded relative to the unfiltered dataset.

| Dataset | All variants | PASS variants | Variants removed | % removed |
|---|---|---|---|---|
| **Single (total)** | 1247235 | 573943 | 673292 | **54.0%** |
| └─ Single-only | 526764 | 185356 | 341408 | **64.8%** |
| **Joint (total)** | 746011 | 448154 | 297857 | **39.9%** |
| └─ Joint-only | 25540 | 59567 | — (increase) | — |

The single-sample dataset decreased from 1,247,235 to 573,943 variants after filtering, corresponding to a loss of 54.0% of calls. In contrast, the joint-genotyped dataset decreased from 746,011 to 448,154 variants, representing a reduction of 39.9%. Notably, variants exclusive to the single-sample analysis exhibited the highest proportional reduction, with 64.8% being removed after filtering, indicating that a large fraction of these calls correspond to low-confidence variants. These results demonstrate that joint genotyping produces a more conservative and higher-quality variant set, whereas single-sample calling generates a larger number of variants but includes a higher proportion of low-confidence calls that are subsequently removed by quality filtering.

#### Table S2. Summary statistics of variant counts per sample across cohorts, variant classes, and genotyping strategies

**Table S2.** Summary statistics of variant counts per sample across sequencing data types (WES or mtDNA), variant classes (SNVs or INDELs), cohorts (phs001172 or 1000 Genomes Project), and genotyping strategies (single-sample or joint calling). For each combination, the table reports the number of samples analyzed and the distribution of variant counts per VCF file (minimum, first quartile (Q1), median, third quartile (Q3), and maximum). These statistics were used to assess cohort comparability and the impact of genotyping strategy on variant discovery.

| Sequencing Data | Genotyping Strategy | Variant Type | Cohort | # samples | Min # variants in vcf | Q1 | Median | Q3 | Max # variants in vcf |
|---|---|---|---|---|---|---|---|---|---|
| WES | Single-sample | SNVs | 1000G | 503 | 15984 | 19714 | 22338 | 23515 | 30704 |
| WES | Joint | SNVs | 1000G | 503 | 14694 | 19901 | 22444 | 23759 | 30890 |
| WES | Single-sample | INDELs | 1000G | 503 | 415 | 514 | 578 | 631 | 891 |
| WES | Joint | INDELs | 1000G | 503 | 464 | 655 | 720 | 803 | 1165 |
| mtDNA | NA | SNVs | 1000G | 493 | 1 | 3313 | 3911 | 4143 | 4725 |
| mtDNA | NA | INDELs | 1000G | 493 | 0 | 40 | 43 | 52 | 84 |
| WES | Single-sample | SNVs | phs001172 | 608 | 15509 | 21911 | 22502 | 23087 | 64401* |
| WES | Joint | SNVs | phs001172 | 608 | 15632 | 21886 | 22627 | 23334 | 38238* |
| WES | Single-sample | INDELs | phs001172 | 608 | 261 | 501 | 594 | 669 | 2309 |
| WES | Joint | INDELs | phs001172 | 608 | 337 | 631 | 759 | 856 | 1624 |
| mtDNA | NA | SNVs | phs001172 | 553 | 1 | 1898 | 2866 | 4214 | 4824 |
| mtDNA | NA | INDELs | phs001172 | 553 | 0 | 26 | 40 | 49 | 113 |

* The maximum values observed (30k~64k variants) correspond to a small number of outlier samples (Sample SRR5090246), whereas the median variant counts remain highly consistent across cohorts.
Note: mtDNA sample numbers differ from the initial cohort, as a subset of samples did not meet the MToolBox coverage threshold required for reliable variant calling (10/503 for 1000G; 55/608 for dbGaP).

For WES single-sample SNVs, both cohorts exhibited nearly identical median values (22,502 for phs001172 and 22,338 for 1000 Genomes (1000G)), with highly overlapping interquartile ranges, indicating comparable variant discovery rates and sequencing depth. A similar pattern was observed for WES INDELs, with medians of 594 and 578 variants per sample for phs001172 and 1000 Genomes, respectively.

Mitochondrial variant counts also showed broadly comparable distributions between cohorts, although modest differences were observed, particularly for SNVs, where the median number of variants was slightly higher in the 1000 Genomes cohort (3,911) than in phs001172 (2,866). These differences are likely attributable to technical factors such as variable mitochondrial coverage in WES capture protocols and differences in sequencing depth rather than biological divergence.

We further evaluated the impact of genotyping strategy on variant discovery by comparing single-sample and joint-calling approaches. For WES SNVs, the median number of variants per sample obtained by joint calling (22,444 for 1000G and 22,627 for phs001172) was highly consistent with those observed in the single-sample analyses of the respective cohorts. A similar trend was observed for INDELs, with joint-calling medians (720 for 1000G and 759 for phs001172) falling within the interquartile ranges of both single-sample datasets.

Overall, the similarity in variant count distributions across cohorts and analysis strategies supports the suitability of the phs001172 and 1000 Genomes datasets for joint downstream analyses.
